# Intraspecific Polymorphism for the Presence and Absence of Entire Mitochondrial Chromosomes

**DOI:** 10.1101/313544

**Authors:** Zhiqiang Wu, Daniel B. Sloan

**Affiliations:** Department of Biology, Colorado State University, Fort Collins, CO 8052

**Keywords:** multichromosomal mitochondrial genome, next-generation sequencing, phylogeography, *Silene noctiflora*

## Abstract

Although mitochondrial genomes are typically thought of as single circular molecules, these genomes are fragmented into multiple chromosomes in many eukaryotes, raising intriguing questions about inheritance and (in)stability of mtDNA in such systems. A previous comparison of mitochondrial genomes from two different individuals of the angiosperm species *Silene noctiflora* found variation in the presence of entire mitochondrial chromosomes. Here, we expand on this work with a geographically diverse sampling of 25 *S. noctiflora* populations. We also included the closely related species *S. turkestanica* and *S. undulata*, with the latter exhibiting a surprising phylogenetic placement nested within the diversity of *S. noctiflora* mitochondrial haplotypes. Using a combination of deep sequencing and PCR-based screening for the presence of 22 different mitochondrial chromosomes, we found extensive variation in the complement of chromosomes across individuals. Much of this variation could be attributed to recent chromosome loss events. Despite the fragmented structures of these mitochondrial genomes and the evidence for occasional biparental inheritance in other *Silene* species, we did not find any indication of recombination between distinct mitochondrial haplotypes either within or among mitochondrial chromosomes, which may reflect the extreme paucity of nucleotide sequence polymorphism and/or the high selfing rate in this species. These results suggest that the massively expanded and fragmented mitochondrial genomes of *S. noctiflora* may have entered a phase of genome reduction in which they are losing entire chromosomes at a rapid rate.

## Introduction

Mitochondrial genome architecture is remarkably diverse (Gray *et al.*, 1999; Smith and Keeling, 2015). Most mitochondrial genomes are represented by a single chromosome, which in some cases can even retain much of its ancestral bacterial-like architecture (Lang *et al.*, 1997; Burger *et al.*, 2013). But many independent eukaryotic lineages have evolved complex multichromosomal structures (Lukes *et al.*, 2005; Shao *et al.*, 2009; Vlcek *et al.*, 2011; Smith *et al.*, 2012). The evolutionary consequences of dividing a genome into multiple chromosomes are not well understood and pose fundamental questions about inheritance and genome stability.

Flowering plants are particularly extreme with respect to their diverse and unusual mitochondrial DNA (mtDNA) structures (Mower *et al.*, 2012). Angiosperm mitochondrial genomes are very large and contain recombinationally active repeat sequences, resulting in complex and dynamic structures *in vivo*, but they can typically be mapped as a “master circle” structure (Sloan, 2013). In contrast, multichromosomal mitochondrial genomes have been identified in five independent angiosperm genera: *Amborella* (Rice *et al.*, 2013), *Cucumis* (Alverson *et al.*, 2011), *Lophophytum* (Sanchez-Puerta *et al.*, 2017), *Saccharum* (Shearman *et al.*, 2016), and *Silene* (Sloan *et al.*, 2012). The most dramatic examples have been found in certain *Silene* species in which the mitochondrial genome has expanded enormously (up to 11 Mb in size) and been fragmented into dozens of circular-mapping chromosomes (Sloan *et al.*, 2012).

In a recent comparison of sequenced mitochondrial genomes from two different populations of *Silene noctiflora* (OSR and BRP), we found that the two genomes were highly similar in sequence and structure, with the one major exception that they differed in their numbers of chromosomes (59 vs. 63) (Wu, Cuthbert, *et al.*, 2015). Each genome contained many unique chromosomes that were absent altogether in the other, suggesting that the dominant mode of molecular evolution in this system is acting at the level of entire chromosomes. However, we could not determine whether variation in the presence of any given chromosome was the result of a recent gain in one lineage or a recent loss in the other. One hypothesis is that the differences in chromosome content are the result of an ongoing process of simple segregational loss during mitochondrial division and cell division that followed an ancestral expansion and fragmentation of the mitochondrial genome. Although most of the “missing” chromosomes contain at least some transcribed regions (Wu, Stone, *et al.*, 2015), they generally have no identifiable genes and are populated by large amounts of non-coding sequence of unrecognizable origin. Therefore, they may experience little or no functional constraint that would prevent such losses.

The fragmentation of the *S. noctiflora* mitochondrial genome raises additional questions about whether it might facilitate independent assortment of chromosomes as a mechanism that generates novel combinations of alleles (Rand, 2009; Wu, Cuthbert, *et al.*, 2015). Although maternal inheritance is the predominant mode of mitochondrial transmission in angiosperms, evidence of low frequency paternal “leakage” of mtDNA has been observed in both natural populations and controlled crosses in many species, including some within the genus *Silene* (McCauley, 2013). Moreover, mitochondria readily and regularly fuse with each other, allowing for intermixing of different copies of the mitochondrial genome (Arimura and Tsutsumi, 2005; Chan, 2006; Segui-Simarro *et al.*, 2008), and crossovers between repeated/homologous sequences frequently occur in plant mtDNA (Arrieta-Montiel and Mackenzie, 2011). Therefore, many of the ingredients for sexual-like inheritance and recombination may already be in place for plant mitochondrial genomes (Stadler and Delph, 2002; Touzet and Delph, 2009; McCauley, 2013; Delph and Montgomery, 2014; Levsen *et al.*, 2016). For simplicity, we will refer to this process as “sexual recombination” to highlight the potential to bring together distinct mitochondrial haplotypes through biparental inheritance and to generate novel combinations of alleles, even though the process does not include meiosis and meet some formal definitions of “sex”.

In this study, we use collections from widespread natural populations of *S. noctiflora* and some of its closest relatives to describe its diversity in mitochondrial genome content and to address the following three questions: 1) How much variation is there in the presence/absence of entire mitochondrial chromosomes? 2) To what extent is that variation the result of recent gains vs. losses of chromosomes? 3) Is there evidence of a history of sexual recombination within and among the mitochondrial chromosomes?

## Materials and Methods

### *Silene* sampling and DNA extraction

*Silene noctiflora* is native to Eurasia but has been widely introduced across the globe as a weedy species (McNeil 1970). To identify variation in mitochondrial genome content across the species range of *S. noctiflora,* we obtained seeds from 25 geographically dispersed populations from Europe and North America (Table S1). In addition, we obtained seeds from a single collection of *S. undulata* (=*S. capensis*), a South African species that has been identified as a close relative of *S. noctiflora* (Havird *et al.*, 2017) (B. Oxelman, pers. comm.). Seeds were germinated on soil (Fafard 2SV Mix supplemented with vermiculite and perlite) in SC7 Cone-tainers (Stuewe and Sons) in February 2014. Plants were grown for two months with regular watering and fertilizer treatments under supplemental lighting (16-hr/8-hr light/dark cycle) in the Colorado State University greenhouse. To test for variation on a more local geographical scale, we also sampled leaf tissue from 19 *S. noctiflora* individuals collected from three sites that are within 10 km of each other in a metapopulation near Mountain Lake Biological Station in southwestern Virginia that is the subject of an annual *Silene* census (Fields and Taylor, 2014) (Table S2). Total cellular DNA was extracted from rosette leaf tissue with a Qiagen Plant DNeasy Kit following the manufacturer’s protocol. We also used previously extracted DNA from *S. turkestanica*, which is a close relative of *S. noctiflora* (Sloan *et al.*, 2009; Rautenberg *et al.*, 2012). To generate sufficient template material from the herbarium-derived *S. turkestanica* DNA sample, we performed whole-genome amplification with a Qiagen Repli-G Mini Kit.

### Chromosome sampling and PCR presence/absence screening

To assess the variation in the presence/absence of specific mitochondrial chromosomes in *Silene noctiflora*, we chose a sample of 22 chromosomes, which were divided into four different groups based on our previous comparison of the mitochondrial genomes from the *S. noctiflora* OSR and BRP populations: 1) the five chromosomes with the highest level of nucleotide sequence divergence, 2) five chromosomes that were present in both OSR and BRP but did not contain any identifiable genes, 3) six chromosomes found in OSR but not in BRP, and 4) six chromosomes that were found BRP but not is OSR. For each of the 22 sampled chromosomes, three distantly spaced PCR primer pairs were designed using Primer3 (Untergasser *et al.*, 2012) (Table S3).

All extracted DNA samples were quantified with a Nanodrop 2000 UV spectrophotometer (Thermo-Fisher Scientific) and diluted to a concentration of 0.5 ng/μl. For each PCR amplification, two replicates were performed to verify consistent determination of marker presence/absence. PCRs were performed in a Bio-Rad C1000 Touch Thermal Cycler in 10 μl reaction volumes containing 1 ng template DNA, 0.2 μM concentration of each primer, 0.1 mM concentration of each dNTP, 1 μl 10× buffer, and 0.1 U Paq5000 DNA polymerase (Agilent Technologies). Amplification was achieved using 3 min of initial denaturation at 94 °C, 38 cycles of 15 s at 94 °C, 15 s annealing at 54 °C, and 30 s extension at 72 °C, followed by a final 5-min incubation at 72 °C. The reactions were screened for the presence/absence of an amplified fragment with the expected size on a 1.5% agarose gel using a 1 Kb plus Ladder (Thermo-Fisher Scientific) as a molecular size standard.

### Illumina genomic DNA sequencing and read mapping

To get a more detailed assessment of the presence/absence of mitochondrial genome content, we used Illumina sequencing of total cellular DNA from individuals from two different populations (KEW 22121 and OPL). Each DNA sample was used for Illumina library construction and sequenced on an Illumina HiSeq 2500 platform (1×76 bp reads for KEW 22121 and 2×151 bp reads for OPL). Library construction and sequencing were performed at the Yale Center for Genome Analysis. The resulting sequencing reads were submitted to the NCBI Sequence Read Archive (SRP140576). Read quality was assessed using FastQC version 0.10.1 (http://www.bioinformatics.babraham.ac.uk/projects/fastqc/), and adapter and low-quality sequences were trimmed using either Cutadapt v1.3 (Martin, 2011) or Trimmomatic version 0.32 (Bolger *et al.*, 2014). The filtered and trimmed reads were then mapped to previously published reference mitochondrial genomes from *S. noctiflora* OSR and BRP (Sloan *et al.*, 2012; Wu, Cuthbert, *et al.*, 2015) using BWA v 0.7.12 (Li and Durbin, 2009) under default parameters. Samtools v1.3 (Li *et al.*, 2009) was used to calculate read-depth in a 1-kb sliding window analysis across the length of each chromosome. Results were visualized using the ggplot2 package (http://ggplot2.org/) in R v3.2.4 (www.r-project.org).

### Sampling of mitochondrial loci and Sanger sequencing

To infer the mitochondrial genealogy of the sampled *S. noctiflora* populations, we designed nine primer pairs (Table S4) targeting three coding regions (*cox1*, *mttB*, and *nad2*) and six non-coding regions from the mitochondrial genome and used them to perform PCR and Sanger sequencing. Previous comparisons of whole mitochondrial genomes from two populations (OSR and BRP) found extremely low rates of sequence polymorphisms. Therefore, the above markers were chosen to include regions known contain single-nucleotide polymorphisms (SNPs) (Wu, Cuthbert, *et al.*, 2015). These markers were used for sequencing all of the 25 *S. noctiflora* populations as well as *S. undulata* and *S. turkestanica* (but only five of the nine markers could be amplified for *S. turkestanica*). PCR amplification was conducted as described above, and the resulting PCR products were purified and sent to University of Chicago Comprehensive Cancer Center DNA Sequencing Facility for Sanger sequencing.

These data were combined with sequence data from four additional mitochondrial coding regions (*atp1*, *atp9*, *cox3*, and *nad9*) for which there has been thorough sampling across the genus *Silene*, including representatives of *S. noctiflora* and *S. turkestanica* (Sloan *et al.*, 2009; Rautenberg *et al.*, 2012). We did not attempt to amplify and sequence these loci in all *S. noctiflora* populations, but we extracted the corresponding sequences from the published mitochondrial genomes of OSR and BRP (Sloan *et al.*, 2012; Wu, Cuthbert, *et al.*, 2015). We also mapped the Illumina sequencing data (see above) to the *S. noctiflora* OSR mitochondrial genome using the CLC Genomics Workbench v7.5.1 (www.clcbio.com) to determine the corresponding sequences for KEW 22121 and OPL. The sequences of *atp1*, *atp9*, *cox3*, and *nad9* were obtained for *S. undulata* in a similar fashion by mapping previously obtained RNA-seq reads (Havird *et al.*, 2017). To avoid misinterpreting sequence changes introduced by RNA editing, all variants associated with known editing sites in *Silene* (Sloan *et al.*, 2010) were removed. Finally, sequence data for all seven coding loci (but not the six non-coding loci) were obtained from published genome assemblies for the outgroups *S. latifolia*, *S. vulgaris*, and *Dianthus caryophyllus* (Sloan *et al.*, 2012; Yagi *et al.*, 2014).

### Sequence alignment, phylogenetic analysis, and ancestral state reconstruction

Individual genes were aligned separately and concatenated using BioEdit (Hall, 1999) and ClustalX v2.1 (Larkin *et al.*, 2007). Phylogenetic trees were inferred by maximum likelihood (ML) implemented with PHYML Version 2.4.5 (Guindon and Gascuel, 2003). The ML analysis was performed with 1000 bootstrap replicates under the GTR model of nucleotide substitution with a BIONJ tree as a starting tree. To infer the history of chromosome gain/loss in *S. noctiflora*, a parsimony criterion was applied to reconstruct the ancestral state for the presence/absence of each individual chromosome using Mesquite v3.0 and the tree topology inferred from the ML analysis described above (Maddison and Maddison, 2006).

### Tests of sexual recombination

To test for a history of sexual recombination among mitochondrial loci/chromosomes, variants identified from the nine amplified markers were subjected to the “four-gamete” test (Hudson and Kaplan, 1985; McCauley, 2013). DnaSP version 5.10.1 (http://www.ub.edu/dnasp/) and the RminCutter.pl script (https://github.com/RILAB/rmin_cut/blob/master/RminCutter.pl) were used to identify the minimum number of recombination events required to fit the data.

### Quantitative PCR Validation

Unexpectedly, one chromosome (OSR chromosome 46) that had previously been found to be absent from the BRP mitochondrial genome (Wu, Cuthbert, *et al.*, 2015), produced consistent amplification across all 25 *S. noctiflora* populations, including BRP. To assess whether this chromosome might be present at lower copy number in some individuals, allowing it to have escaped detection in earlier genome sequencing efforts, we performed quantitative PCR (qPCR) using individuals from each of six different *S. noctiflora* populations (BRP, OSR, MH-L, BWT, KEW 12991 and KEW 36186) and two technical replicates. We also analyzed the relative copy number of chromosome 37, which is present in both BRP and OSR, to serve as a control. For each chromosome, three pairs of qPCR primers were designed (Table S5). We also selected two mitochondrial protein-coding genes (*cox1* and *nad2*) to serve as reference markers. All qPCR amplifications were performed in 10-μL volumes with Bio-Rad SsoAdvanced SYBR Green 2× Supermix, 0.2 μM concentration of each primer, and 1 ng of template DNA. Amplification was performed on a Bio-Rad CFX96 Touch Real-Time PCR Detection System (Bio-Rad) with an initial 3-min incubation at 95 °C and 40 cycles of 3 s at 95 °C and 20 s at 60 °C, followed by a melt curve analysis. Copy number of the two chromosomes relative to the reference genes were estimated for each sample with the geNorm method (Vandesompele *et al.*, 2002).

## Results

### Mitochondrial Genealogy

In order to reconstruct the history of mitochondrial chromosome gain and loss in this lineage, we first inferred the mitochondrial genealogy using sequence data from 13 markers. This analysis provided evidence for four distinct, well-supported mitochondrial lineages within *S. noctiflora* (Figure 1). Two of these lineages correspond to the BRP and OSR backgrounds that have already been thoroughly characterized based on complete mtDNA sequences (Wu, Cuthbert, *et al.*, 2015). Each of these two lineages is represented by multiple populations in our dataset (eight in the BRP-like group and 15 in the OSR-like group). Two additional lineages were also detected, each represented by only a single sample (KEW 1672 and KEW 22121). Despite the broad geographical sampling across Europe and North America (Table S1), the overall level of intraspecific polymorphism was extremely low, and no sequence variants were detected among populations within the BRP- or OSR-like groups, which is consistent with the general lack of mitochondrial sequence diversity in previous studies of *S. noctiflora* (Sloan *et al.*, 2012; Wu, Cuthbert, *et al.*, 2015). This mitochondrial data set confirmed the close relationships between *S. noctiflora*, *S. turkestanica*, and *S. undulata* relative to the rest of the genus *Silene* (Sloan *et al.*, 2009; Rautenberg *et al.*, 2012; Havird *et al.*, 2017). Unexpectedly, phylogenetic analysis placed the *S. undulata* mitochondrial haplotype as nested within the small amount of observed diversity in *S. noctiflora*. It formed a well-supported group with the *S. noctiflora* BRP-like and KEW 1672 lineages to the exclusion of the KEW 22121 and OSR-like lineages (Figure 1).

**Figure 1.**
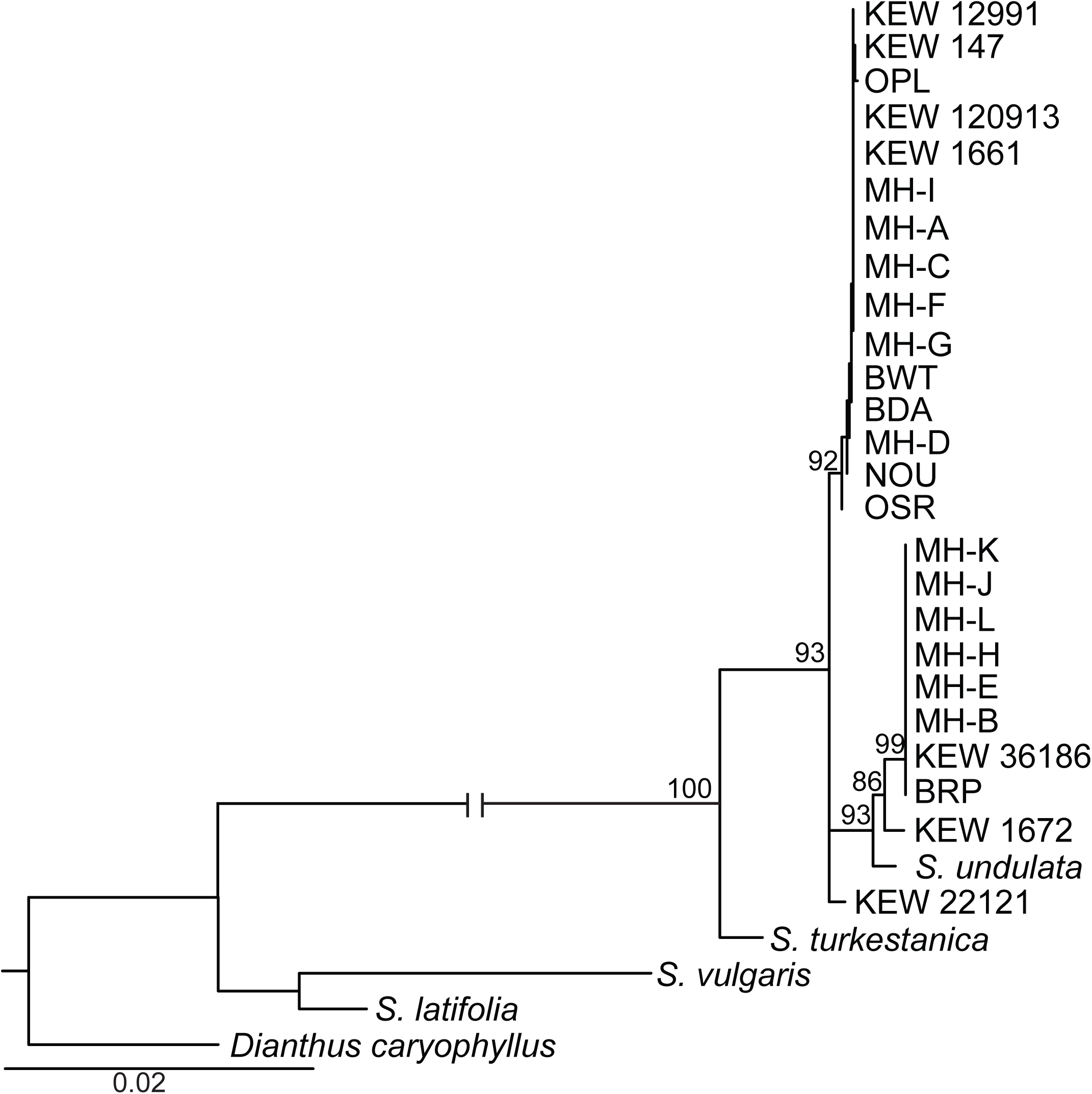
Phylogenetic relationships among 25 *S. noctiflora* samples and closely related species based on 13 mitochondrial markers. Supporting bootstrap values greater than 80 are shown for each node.

### Variation in Mitochondrial Chromosome Presence/Absence

PCR-based screening for a sample of 22 mitochondrial chromosomes revealed substantial variation in chromosome presence/absence among populations of *S. noctiflora* and related species (Figure 2), especially when juxtaposed with the extremely low levels of nucleotide sequence divergence. Among the *S. noctiflora* populations, the patterns of chromosome presence/absence mirrored the sequence-based analysis and clustered into the same four lineages, with *S. undulata* exhibiting a fifth distinct pattern. Although the 15 *S. noctiflora* OSR-like mitochondrial haplotypes were all identical across the sequenced mitochondrial markers (see above), there was variation in chromosome presence/absence within this group, with pairs of samples differing in the presence/absence of up to four chromosomes (Figure 2). In contrast, all eight BRP-like samples had identical presence/absence profiles. Only four of the sampled mitochondrial chromosomes were detectable at all in *S. turkestanica*, and none of those chromosomes produced positive amplification for all three markers, indicating that mitochondrial genome content and structure is highly divergent in *S. turkestanica* relative to *S. noctiflora/undulata*. Using a simple parsimony criterion and the genealogy inferred from mitochondrial sequence data, we inferred ancestral states for the presence/absence of each sampled chromosomes (Figure 3).

**Figure 2.**
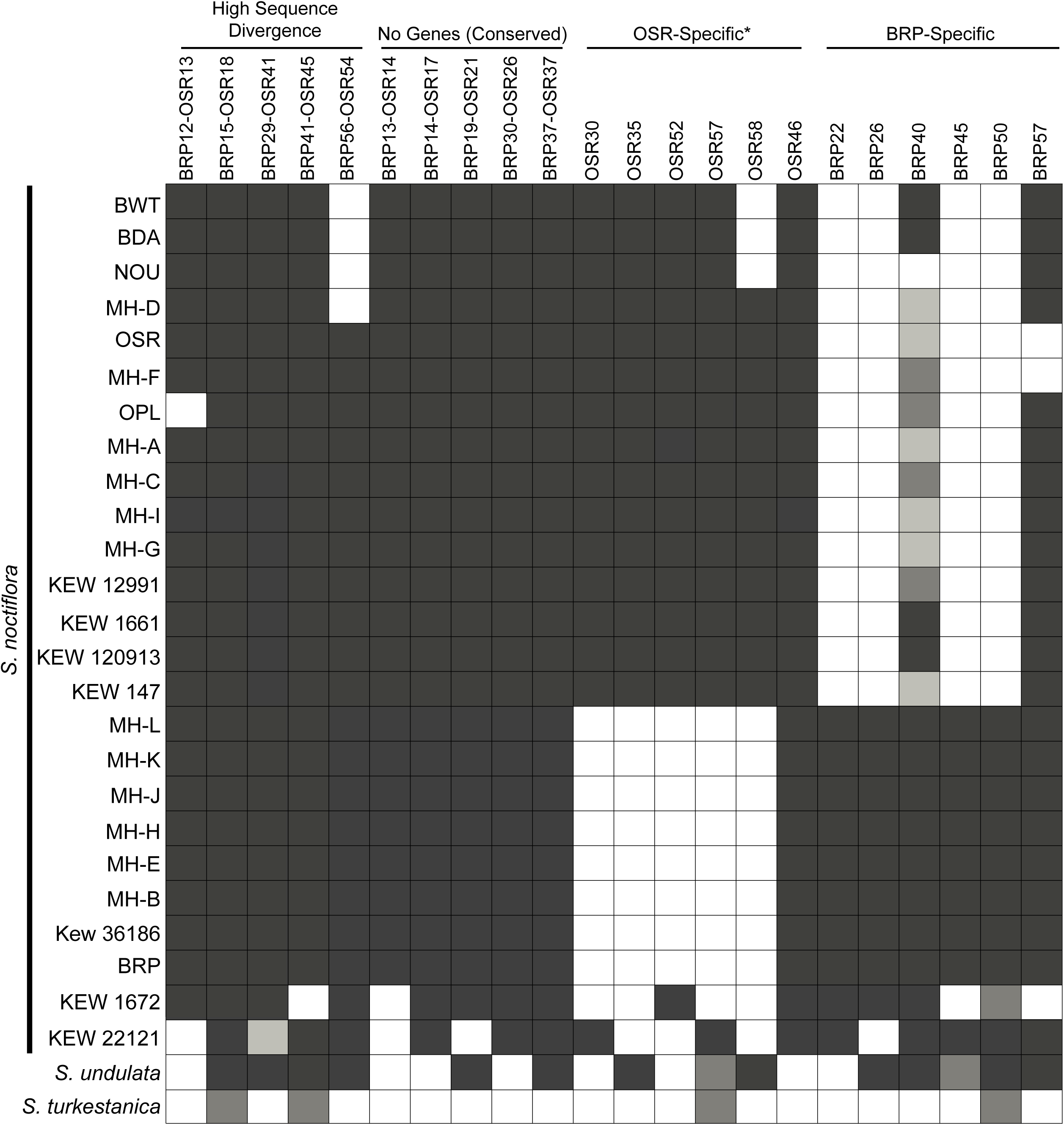
Presence/absence survey of 22 mitochondrial chromosomes from 25 different populations of *S. noctiflora* (Table S1) and two closely related species. For each chromosome, presence/absence was assessed with three different PCR markers in each chromosome. Dark gray shading indicates positive detection for all three markers; medium gray shading indicates positive detection for two of three markers; light gray shading indicates positive detection for only one of three markers. For three chromosomes, the analysis was performed with only two markers (BRP41-OSR45, OSR58, and BRP57). In these cases, dark gray shading indicates detection of both markers. The chromosomes are divided into four categories as described in the Methods. Note that one of the “OSR-specific” chromosomes (OSR46) was found to be present at low levels in BRP and other samples from that group (see Results).

**Figure 3.**
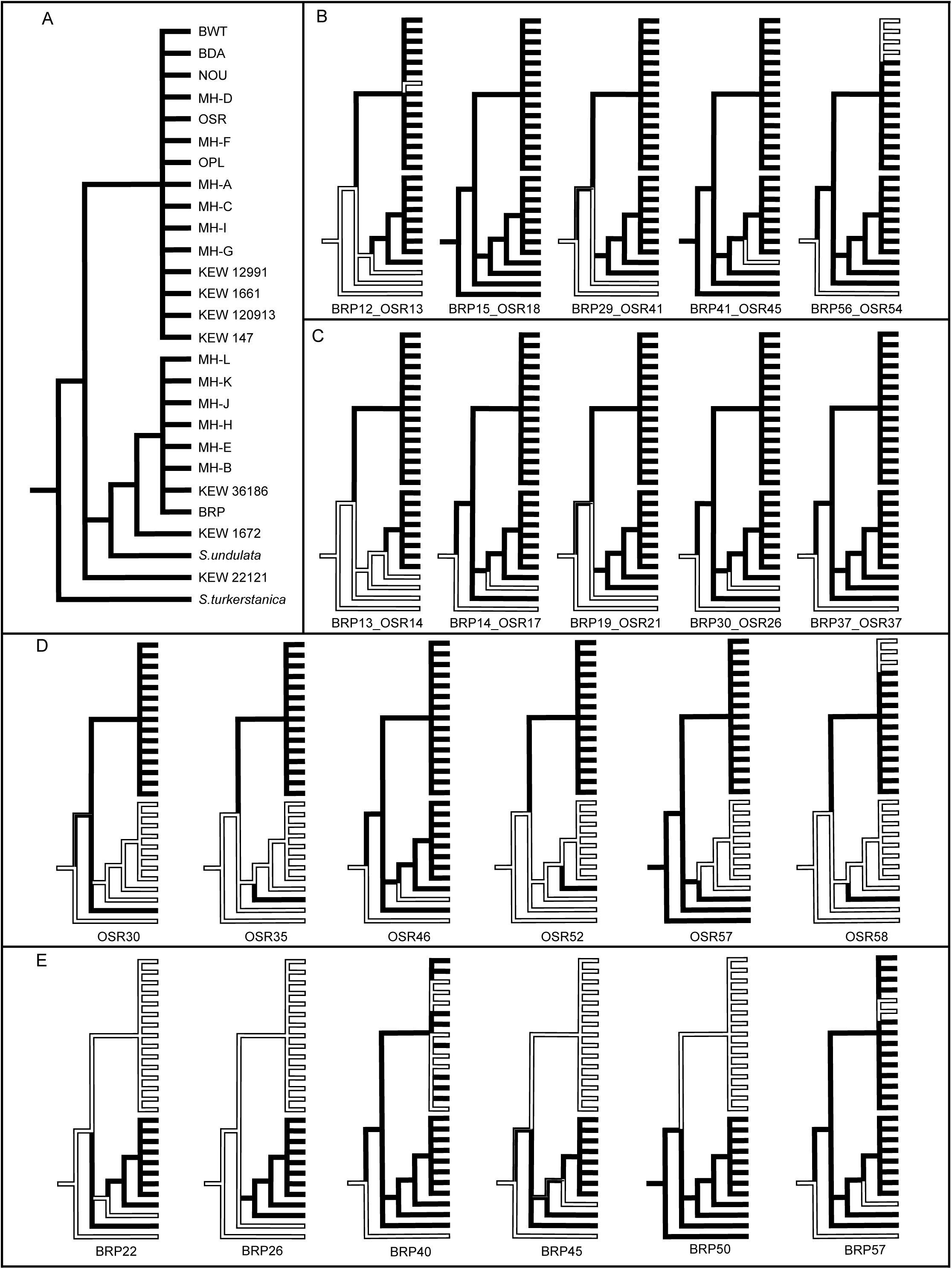
Parsimony-based reconstruction of ancestral States for the presence (black) or absence (white) of each of the 22 sampled chromosomes cross 25 *S. noctiflora* samples and two closely related species. A) The constraint topology used for the analysis based on sequence data from 13 mitochondrial markers (see Figure 1); The remaining panels show the presence/absence states for (B) the five sampled chromosomes with high sequence divergence, (C) the five sampled chromosomes that conserved between BRP and OSR and have no genes, (D) the six sampled “OSR-specific” chromosomes, and (E) the six sampled “BRP-specific” chromosomes.

To characterize variation at a finer level of geographical resolution, we analyzed seeds from five to seven different individuals from each of three sites (< 10 km apart) within a *S. noctiflora* metapopulation in southwestern Virginia near Mountain Lake Biological Station (Table S2). We found that all sampled individuals within each site had the exact same pattern of presence/absence for a sample of 10 chromosomes but that the three sites all had different patterns from each other (Figure S1). The three-observed presence/absence patterns across these sites were consistent with those observed for the BRP-like, OSR-like, and KEW 1672 groups that were analyzed for a larger number of chromosomes (Figure 2). Therefore, multiple mitochondrial haplotypes occur at nearby sites in a metapopulation within the introduced range of *S. noctiflora*, but we did not detect evidence of variation in chromosome presence/absence at the finest scale of within-site sampling.

Our inference of chromosome presence/absence was based on three independent PCR markers on each chromosome. In general, each set of three markers produced consistent results, but there were some cases in which only two of the three markers supported presence or absence (Figure 2), raising questions about changes that have affected partial chromosomes. To address variation at this scale, we mapped Illumina sequencing reads derived from total-cellular genomic DNA from *S. noctiflora* KEW 22121 and OPL samples (Figures 4, S2-S4). Four themes emerged from this analysis. First, the mapping results confirmed predictions for the PCR-based marker analysis (e.g., the lack of coverage from KEW 22121 sequencing for BRP chromosomes 12, 13, 19, and 26; Figure 4). Second, many “missing” chromosomes showed no coverage across their entire length with the exception of very short scattered sequences that presumably reflect the short repeats that are shared with other chromosomes (again, see chromosomes 12, 13, 19, and 26 in Figure 4). Third, most of the other chromosomes had consistent coverage across their full length, although the observed coverage level often differed to some extent among chromosomes (e.g. chromosome 45 vs. chromosome 46 in Figure 4). Finally, a smaller subset of chromosomes exhibited major fluctuations in coverage within the chromosome. Large regions with no coverage (e.g. chromosome 29 in Figure 4) suggest partial chromosome absence, and large regions in which coverage abruptly jumped to higher levels suggest sequence duplications and variation in the number of copies of large repeats within the genome.

**Figure 4.**
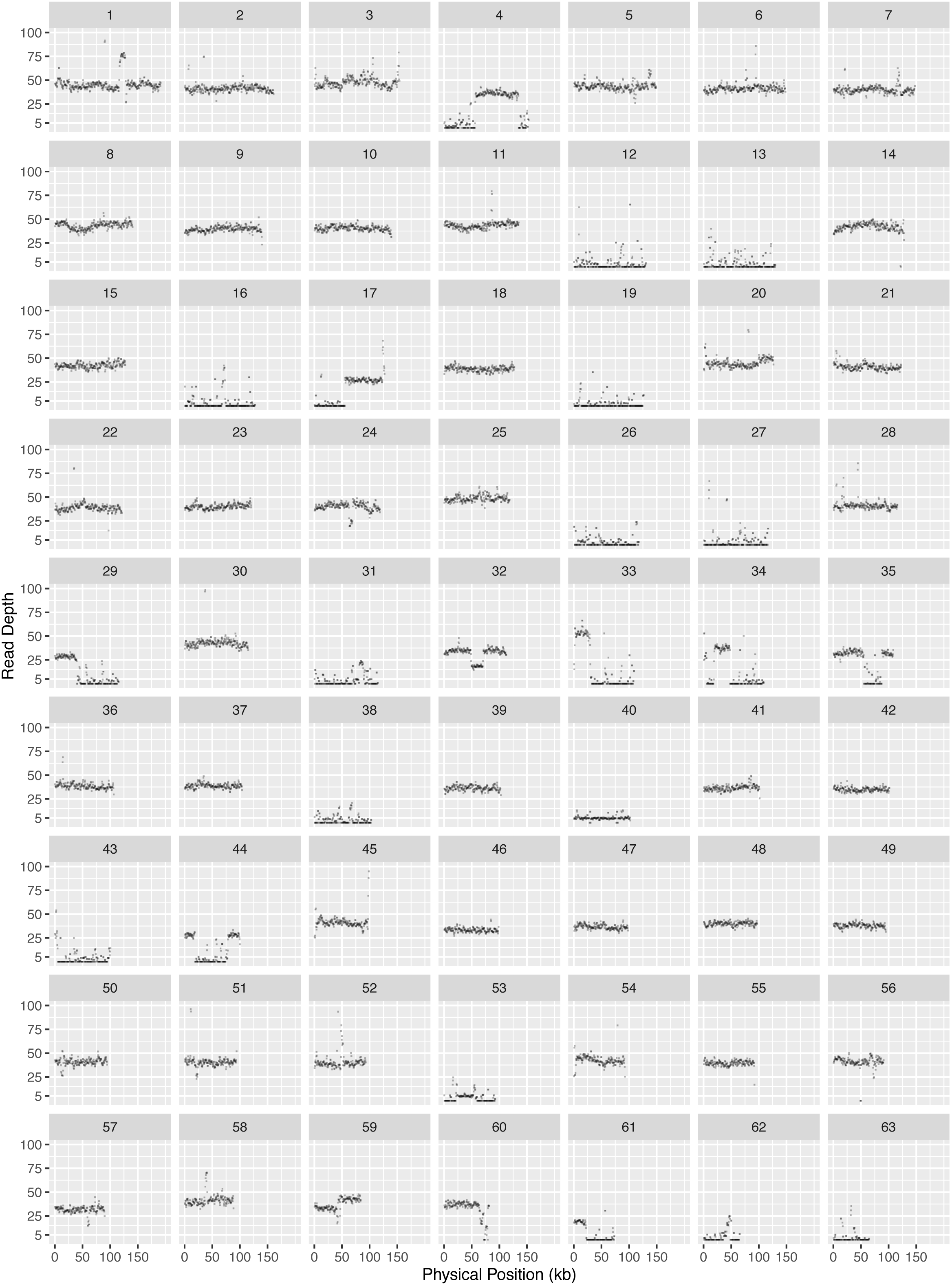
Read depth across the 63 chromosomes of the *S. noctiflora* BRP reference mitochondrial genome based on Illumina sequencing of total-cellular DNA from *S. noctiflora* KEW 22121. Coverage estimates are based on a sliding window with a window size of 1000 bp and a step size of 500 bp. The plot was generated with the ggplot2 library in R.

An unexpected result from our PCR screen was that all three markers for chromosome 46 from the OSR mitochondrial genome were detected in our BRP sample (and all other *S. noctiflora* samples) despite our previous finding that it was absent from the BRP genome assembly (Wu, Cuthbert, *et al.*, 2015). One potential explanation for this discrepancy is that the chromosome was present but at an abundance that was too low to be captured in the earlier BRP sequencing and assembly data set. Consistent with this possibility, our mapping analysis of a different sample (KEW 22121; see above) detected consistent coverage across the full length of the chromosome but at a level much lower than the other chromosomes (see chromosome 46 in Figure S2). We found further support for this interpretation by performing qPCR, which detected the presence of this chromosome in BRP and other BRP-like samples but at a much lower relative copy number (Figure 5). A second potential example of this phenomenon is the chromosome 40 from the BRP mitochondrial genome. This chromosome was not detected in the original OSR genome assembly, but mapping of sequence reads from OPL (a member of the OSR-like group) produced a low by consistent level of coverage across the chromosome (Figure S3). In addition, PCR-based screening produced sporadic amplification across the OSR-like group (often with only very faint bands), including for one of the three markers in OSR (Figure 2).

**Figure 5.**
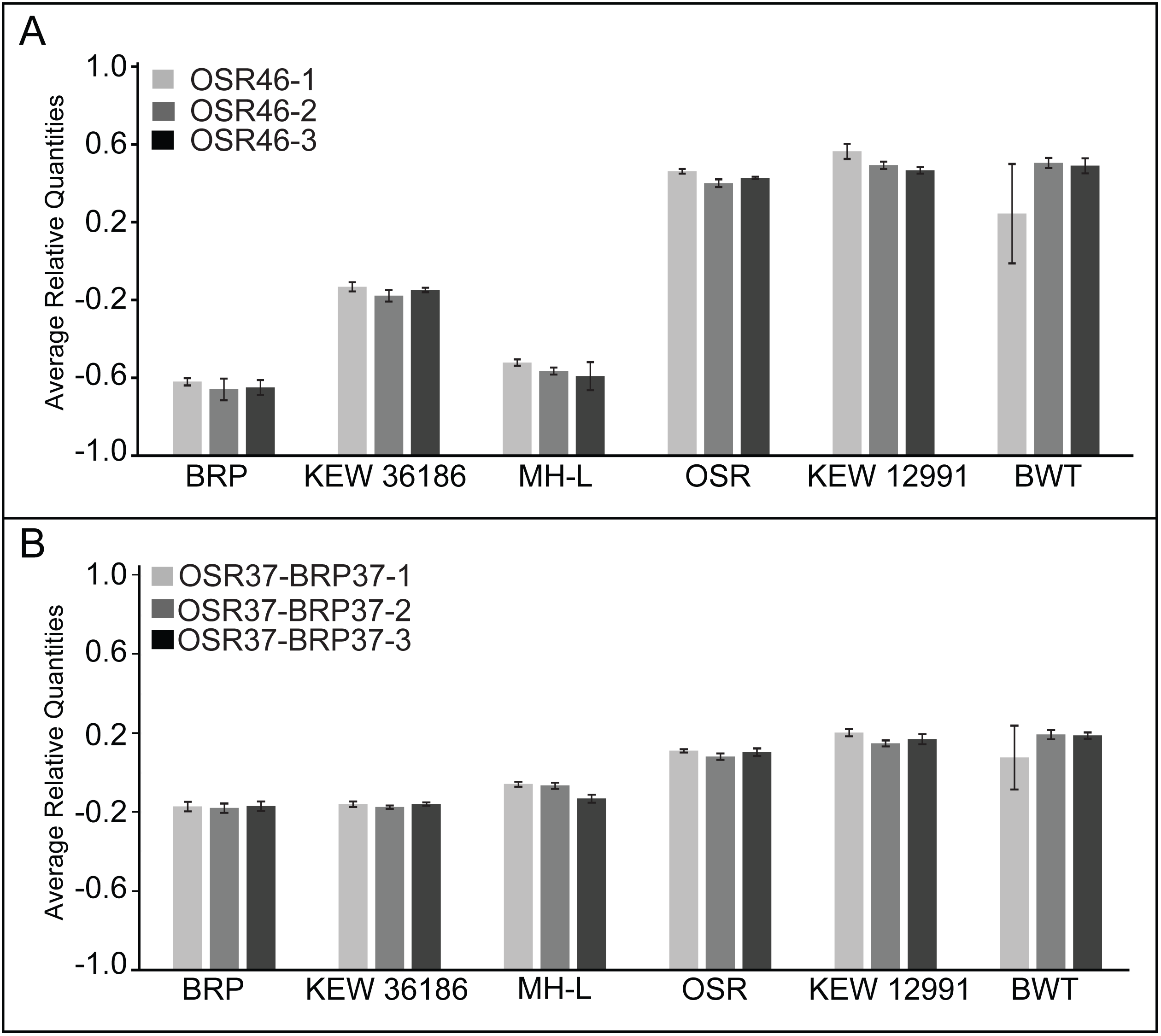
Relative qPCR analysis of copy number variation for two *S. noctiflora* mitochondrial chromosomes OSR46 and OSR37-BRP37 in the six different *S. noctiflora* families. Copy numbers were calculated relative to two mitochondrial genes (nad2 and cox1). For each chromosome, three independent loci were sampled to do the test, and each marker with two technical replicates. Error bars are based on standard deviations between two technical replicates.

Previous analysis of differences in copy number among chromosomes within a sample found that relative abundance does vary but across a relatively narrow range (less than two-fold) (Wu, Cuthbert, *et al.*, 2015). Our observation of the OSR46 chromosome suggest that much wider ranges can occur. However, based on these data, we cannot exclude the alternative interpretation that some of the sampled individuals have lost the mitochondrial chromosome but still harbor an older insertion of this mitochondrial sequence into the nuclear genome (i.e., a *numt*; (Hazkani-Covo *et al.*, 2010)).

### No Evidence of Sexual Recombination among Mitochondrial Markers

We tested for a history of sexual recombination in *S. noctiflora* mitochondrial DNA, using the sequenced markers described above. However, we did not detect any pairs of sites that were present in all possible haplotype combinations (i.e., the four-gamete test). Likewise, analyses with DnaSP and RminCutter.pl, which is based on the *R_M_* statistic ofHudson and Kaplan (Hudson and Kaplan, 1985), also failed to detect any recombination events.

## Discussion

### Gain vs. Loss of Mitochondrial Chromosomes

Our observations confirm and extend previous work showing remarkable intraspecific variation in chromosome content among samples that barely differ in nucleotide sequence (Wu, Cuthbert, *et al.*, 2015). Previous analyses were unable to assess the extent to which these differences in chromosome content reflected recent gains vs. losses of chromosomes. In this study, a simple parsimony reconstruction of ancestral states suggests variation in the timing of gains and losses (Figure 3). These include clear cases of recent chromosome loss, in which mitochondrial markers are broadly shared across the *S. noctiflora* and *S. undulata* samples but are absent from one or a small number of lineages that are deeply nested within the group (e.g., chromosomes BRP12/OSR13, BRP56/OSR54, and BRP57; Figures 2 and 3). One notable example is chromosome 57 in BRP, which is absent from two different highly nested parts of the tree (OSR/MH-F and KEW 1672), providing strong evidence for two independent losses of this chromosome.

This result indicates a high rate of chromosome loss and may be relevant to interpreting the presence/absence patterns of other chromosomes. For example, a simple-parsimony interpretation of chromosomes such as OSR 52 implies two independent gains of the same chromosome (Figure 3). However, unless horizontal transfer of sequence content has occurred, we feel that this inference is less likely than a history in which the chromosome was ancestrally present at the base of the group and then lost independently in multiple lineages. We also note that chromosomes BRP 26 and BRP 40 are only found in the BRP/KEW 1672/*S. undulata* clade (Figure 3), which would imply that those chromosomes were not ancestrally present in *S. noctiflora* and were instead gained more recently at the base of that specific clade. Therefore, it is possible that the history of chromosome gain that shaped the massive, multichromosomal genomes of *S. noctiflora* occurred over a prolonged period that extended past the point when extant lineages within this species began to diversify. However, as above, the simple-parsimony reconstruction may be neglecting the possibility of ancestral presence followed by numerous independent losses. Overall, we conclude that there is strong evidence that recent chromosome losses have played a substantial role in the observed presence/absence variation but the evidence for recent gains is more ambiguous.

There are some important limitations to consider in our analysis to assess the relative contributions of chromosome gain vs. loss in variation among mitochondrial haplotypes. An obvious bias with our PCR-based screen is that the markers were all designed based on previously identified chromosomes. Therefore, they are capable of inferring the history of recent losses of chromosomes known to be shared between BRP and OSR, but they cannot detect recent lineage-specific gains of entirely novel chromosomes that are not found in either of those two genomes. This limitation also applies to our reference-based mapping of sequencing datasets. It is very possible that those samples contain additional chromosomes that are not found in either the OSR or BRP mitochondrial genomes.

### Sexual Recombination in Multichromosomal Mitochondrial Genomes

We and others have speculated that the fragmentation of mitochondrial genomes into multiple chromosomes may facilitate sexual recombination between physically unlinked loci (Rand, 2009; Wu, Cuthbert, *et al.*, 2015), and evidence for sexual recombination has been found in some *Silene* species (Stadler and Delph, 2002; Touzet and Delph, 2009; McCauley, 2013). However, our sampling of mitochondrial markers from multiple chromosomes in this study did not reveal any evidence of sexual recombination. There are multiple factors that might contribute to rarity/absence of mitochondrial recombination in *S. noctiflora* and the lack of power to detect it even if it is occurring. First, because of the very low levels of nucleotide polymorphism, many recombination events might occur between identical sequences and therefore be impossible to detect. Breeding system and modes of mitochondrial inheritance also contribute to the opportunity for biparental sexual reproduction to occur. Although low levels of paternal transmission of mtDNA has been documented in the genus *Silene* (McCauley, 2013), only a small number of crosses have been performed in *S. noctiflora* to track mtDNA inheritance, and they did not identify any paternal leakage (Sloan *et al.*, 2012). Therefore, a lack of biparental mtDNA inheritance in this species may preclude opportunities for sexual reproduction. In addition, *S. noctiflora* has a high rate of self-fertilization (Davis and Delph, 2005), such that even if biparental inheritance of mtDNA occurs, it may rarely bring together two distinct haplotypes.

### Phylogenetic Placement of *Silene undulata*

We included *S. undulata* as a potentially close outgroup to *S. noctiflora* on the recommendation of Bengt Oxelman who recently recognized the relatedness between these two species (pers. comm.). The placement of the *S. undulata* mitochondrial haplotype as nested within the clade of *S. noctiflora* haplotypes was a surprise. These results are especially surprising in light of the distant distribution of *S. undulata* in Southern African compared to the Eurasian distribution of *S. noctiflora.* Despite the extreme similarity in mitochondrial haplotypes, our *S. undulata* sample exhibited morphological features that distinguish it from *S. noctiflora*, including petals that were erose at their apices and unfurled during the daytime. The general lack of polymorphism in *S. noctiflora* is also somewhat perplexing given the species’ broad geographical distribution and historically high mitochondrial mutation rates (Mower *et al.*, 2007; Sloan *et al.*, 2009), likely indicating a very low effective population size and/or recent reversion to low mitochondrial mutation rates (Sloan *et al.*, 2012). Given how frequently cytoplasmic genomes introgress across species boundaries (Rieseberg *et al.*, 1996), it is possible that nuclear genealogies show more separation between *S. noctiflora* and *S. undulata.* However, any interpretation involving introgression between these lineages will have to be reconciled with their geographically disjunct distributions. Overall, this lineage appears to have an unusual and intriguing phylogeographic history that would be well worth systematic investigation.

## Acknowledgements

We thank Lucas Dominguez, Jessica Hurley, and Cody Kalous for their help with PCR screening. We also thank Bengt Oxelman for suggesting *S. undulata* as a likely close relative of *S. noctiflora* and Mark Simmons for helpful discussion about ancestral state reconstruction. We are grateful to the Kew Millennium Seed Bank, Helena Štorchová, Mitra Menon, and especially Michael Hood for providing/collecting seeds. Justin Havird and Joel Sharbrough provided insightful comments on an earlier version of this manuscript. This work was supported by National Institutes of Health (NIGMS R01 GM118046) and start-up funding from Colorado State University.

---

Figure S1.

Presence/absence survey of 10 mitochondrial chromosomes (five OSR-specific and five BRP-specific) for *S. noctiflora* individuals from three sites in a metapopulation from southwestern Virginia.

Figure S2.

Read depth across the 59 chromosomes of the *S. noctiflora* OSR reference mitochondrial genome based on Illumina sequencing of total-cellular DNA from *S. noctiflora* KEW 22121. Coverage estimates are based on a sliding window with a window size of 1000 bp and a step size of 500 bp. The plot was generated with the ggplot2 library in R.

Figure S3.

Read depth across the 63 chromosomes of the *S. noctiflora* BRP reference mitochondrial genome based on Illumina sequencing of total-cellular DNA from *S. noctiflora* OPL. Coverage estimates are based on a sliding window with a window size of 1000 bp and a step size of 500 bp. The plot was generated with the ggplot2 library in R.

Figure S4.

Read depth across the 59 chromosomes of the *S. noctiflora* OSR reference mitochondrial genome based on Illumina sequencing of total-cellular DNA from *S. noctiflora* OPL. Coverage estimates are based on a sliding window with a window size of 1000 bp and a step size of 500 bp. The plot was generated with the ggplot2 library in R.

File S1

Concatenated alignment of 13 mitochondrial sequence markers used for phylogenetic analysis.

